# How does the community structure of dung beetles affect ecosystem functions? Modeling of secondary seed dispersal and seedling establishment

**DOI:** 10.64898/2026.10.10.758268

**Authors:** Ryo Akashi

**Affiliations:** Graduate School of Life Science, Hokkaido University, Sapporo, Hokkaido, Japan

## Abstract

Dung beetles are associated with diverse ecosystem functions. However, their populations are declining globally, raising concerns about the loss of ecosystem functions. Among these services, secondary seed dispersal is crucial for plant recruitment and movement. To reveal the relationship between community composition and ecosystem function, I developed a novel mathematical model simulating seedling establishment that accounts for seed dispersal by two functional groups (tunnelers and rollers), seed predation, and among-seedling competition. The model revealed that when seed predators were absent, rollers contributed more to seedling establishment than tunnelers; consequently, communities with a high proportion of rollers and a low proportion of tunnelers maximized the number of established seedlings. Under high predation pressure, tunneler-abundant communities yielded greater seedling establishment. Under intermediate predation pressure, the relative contributions of the two functional groups depended on the intensity of among-seedling competition: tunnelers enhanced establishment when competition was weak, whereas rollers excelled when competition was strong. This study provides the first mathematical framework for dung beetle-mediated secondary seed dispersal, advancing our understanding of the links between community composition and ecosystem functioning. These findings offer critical insights for anthropogenic biodiversity management, such as the conservation and introduction of dung beetle communities.

## 1 Introduction

Ecological communities consist of diverse species that occupy various functional roles in ecosystems. For instance, in the context of material cycling, species can be classified according to their trophic roles as primary producers, consumers, or decomposers. Functional roles can also be defined by the varied interaction types that link species within ecological networks (Lewinsohn et al. 2006). For example, animals achieve different ecological functions in their interactions with plants, acting as pollinators, herbivores or seed dispersers (Bronstein et al. 2006; Nicholls and Altieri 2013; Stam et al. 2014). In other words, functional distinctions can be made from multiple perspectives, and species contribute multilayered roles and collectively construct the functional diversity of a community. Changes in species composition not only alter taxonomic composition but also affect the functional attributes of communities, potentially influencing ecosystem processes at broader scales (Hooper et al. 2005; Tilman et al. 1997).

Dung beetles (Coleoptera: Scarabaeinae, Aphodiinae and Geotrupidae) are a key group contributing to multiple ecosystem functions such as dung removal, nutrient cycling, plant growth enhancement, and seed dispersal (Nichols et al. 2008). Based on their nesting behavior, dung beetles are conventionally classified into three functional groups: dwellers, tunnelers, and rollers. Dwellers (endocoprids) feed and reproduce directly inside the dung pat; tunnelers (paracoprids) dig a tunnel and bury dung subsurface; and rollers (telecoprids) form a ball of dung and transport it horizontally away from the dung pat (Halffter and Edmonds 1982). Due to their distinct nesting strategies, each functional group contributes differently to ecosystem functions, particularly dung removal and seed dispersal (Slade et al. 2007). Tunneler dung beetles cause a high-density, clumped distribution of seeds because they bury dung directly beneath or adjacent to the dung pat by digging tunnels (Halffter and Edmonds 1982). In contrast, roller dung beetles form dung balls and roll them away from the source, resulting in a lower-density, sparser reallocation of seeds (Halffter and Edmonds 1982; Manns et al. 2020). Therefore, the seed dispersal function provided by dung beetles is tightly linked to their community structure, particularly the relative dominance of tunnelers and rollers.

When assessing seed dispersal functions, previous studies have primarily focused on the number of dispersed seeds (Andresen 2002) or germination rate (Koike et al. 2012); few studies have investigated the consequences of secondary seed dispersal (Andresen and Urrea-Galeano 2022). However, to understand seedling establishment success, it is necessary to consider post-dispersal processes, specifically seedling competition and predation. Seed density is a key factor determining establishment success. High seed density intensifies competition (Harms et al. 2000) but may also facilitate predator avoidance (e.g., through the dilution effect). Thus, clumped seed deposition can have both negative and positive consequences. Empirical studies face limitations in disentangling these opposing effects. It remains unclear whether rollers or tunnelers, each of which generates distinct spatial density distributions of seeds, contribute more effectively to seedling establishment when these density-dependent processes are considered. A theoretical approach accounting for post-dispersal seed survival is required to predict how dung beetle community composition is related to seedling establishment success.

The unresolved relationship between dung beetle communities and plant establishment is also crucial from a conservation perspective because dung beetles are currently impacted by anthropogenic activities, including habitat destruction (Hanski et al. 2007), the impact of veterinary medicines (I. Ishikawa and Iwasa 2020; Wall and Strong 1987), and biological invasions (Filho et al. 2018). Seed dispersal is one of the most critical ecological functions mediated by dung beetles, as it facilitates plant migration and reproduction, potentially influencing primary production. Understanding how anthropogenic pressures alter the ecosystem functions of dung beetles is key to a sustainable society.

Therefore, in this study, I develop a novel mathematical model to simulate seed dispersal by dung beetles and subsequent seedling establishment. This model integrates beetle-mediated seed relocation with two contrasting density-dependent processes affecting seed survival and establishment: negative density-dependent competition among seedlings and positive density-dependent dilution of seed predation risk. Finally, I compare the relative contributions of tunnelers and rollers to seedling establishment and examine how shifts in dung beetle community composition influence the overall number of established seedlings. This framework allows us to predict how changes in beetle community composition will enhance or diminish plant recruitment—a key question for understanding the ecological consequences of dung beetle declines and community shifts.

## 2 Methods

This model focuses on two functional groups of dung beetles that differ markedly in their seed dispersal behaviors: tunnelers, which bury dung and seeds near the dung pat, and rollers, which transport dung balls and embedded seeds over longer distances (Figs. 1a, b), Halffter and Edmonds 1982). For simplicity, the model assumes that the dung produced by a herbivore species contains seeds from only a single plant species. The model tracks seed fates after being deposited by herbivores across three stages: dispersal by dung beetles, mortality caused by seed predators, and seedling establishment under among-plant competition.

**Figure 1:**
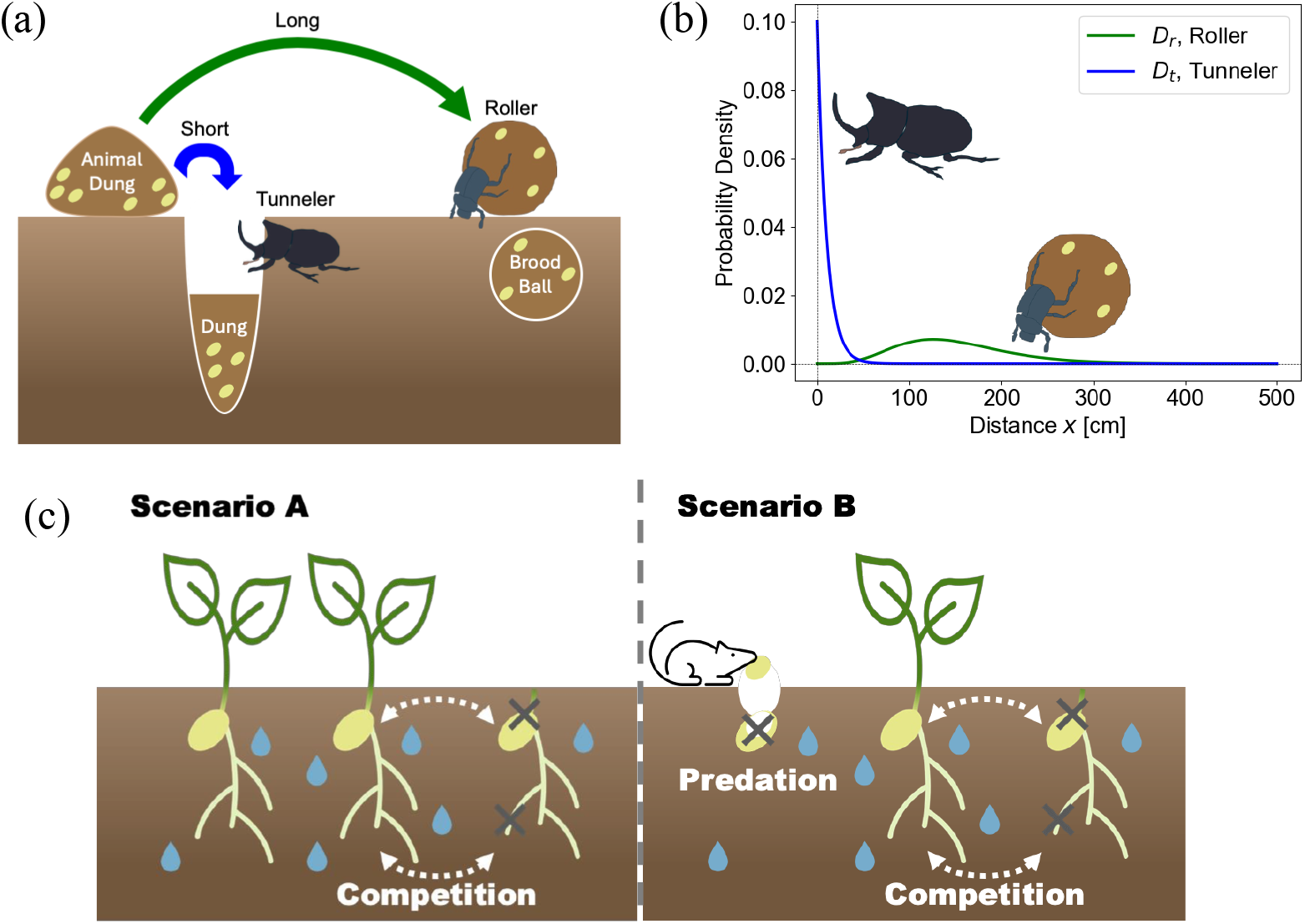
Overview of the situation and scenarios considered in this study. (a)Diagram of the secondary seed dispersal by two types of dung beetles (tunneler and roller). (b) Dispersal distances of the two dung beetle guilds. The probability density distributions of seeds dispersed by tunnelers (blue) and rollers (green). (c) Two scenarios of the post-dispersal processes: Scenario A considers only seedling competition, whereas Scenario B includes both seed predation and seedling competition.

Here, I analyze two scenarios that differ in the post-dispersal processes considered: Scenario A omits seed predation and examines only density-dependent competition, while Scenario B incorporates both density-dependent competition and predation (Fig. 1c). Seed dispersal is a process that prevents the clustering of seeds (Howe and Smallwood 1982). Secondarily dispersed seeds that have experienced endozoochory exhibit low dormancy rates (Razanamandranto et al. 2004) and are thus expected to face among-seedling competition. Therefore, negative density-dependent competition is an essential process to consider when evaluating the consequences of seed dispersal on seedling recruitment (Harms et al. 2000), and the presence or absence of predators represents the environmental conditions where seeds are deposited. Comparing the two scenarios reveals the relationships between community structure and ecosystem functions under different environmental conditions.

### 2.1 Model

#### 2.1.1 Dung beetle community

I assume a dung beetle community consisting of one tunneler and one roller species. The total number of individuals of the dung beetles is fixed at its carrying capacity (*K*). Let *s* denote the proportion tunnelers within this community. The population sizes of tunnelers (*n_t_*) and rollers (*n_r_*) are given by:

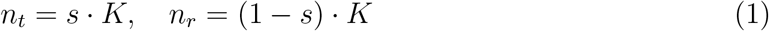

#### 2.1.2 Seed dispersal by dung beetles

For simplicity, we consider one-dimensional seed distribution relocated by dung beetles. I modeled the probability distribution of the density of dispersed seeds with distance, *D_i_*(*x*), defined by the gamma distribution as follows:

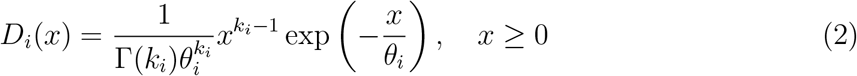

where Γ(*·*) denotes the Gamma function with a shape parameter *k_i_*and a scale parameter *θ_i_*. The subscript *i* denotes the beetle functional group (*i* = *t*: tunneler; *i* = *r*: roller), and the variable *x* represents the distance from the original dung pat (Fig. 1b).

I assume that each beetle functional group consumes dung at a constant per-capita amount: *F_t_* for tunnelers and *F_r_*for rollers. The total dung consumption of the tunnelers and rollers is *F_t_n_t_* and *F_r_n_r_*, respectively. Thus, the proportions of seeds relocated by tunnelers and rollers are given by *F_t_n_t_/*(*F_t_n_t_* + *F_r_n_r_*) and *F_r_n_r_/*(*F_t_n_t_* + *F_r_n_r_*), respectively. I also assume that dung pats are sufficiently isolated from one another, ensuring that the dispersal kernels of seeds from different pats do not overlap. Therefore, the initial local density of seeds that are relocated to a distance *x*, denoted as *N* (0, *x*), is the weighted average of the two dispersal kernels:

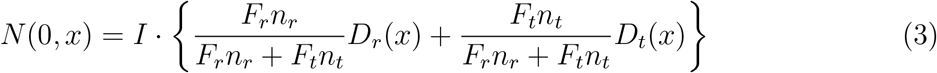

where *I* is the total number of seeds dispersed from a single dung pat.

#### 2.1.3 Seed predation

Next, I model seed predation after seed dispersal, which is considered only in Scenario B. Dispersed seeds may be consumed by seed predators (Janzen 1971). Because the intake rate of seed predators is limited by handling time (Ruscoe et al. 2005), I assume that the predation behavior follows a Type II functional response dependent on the local seed density *N* (Holling 1959). The per capita predation rate at distance *x* and specific time *τ*, denoted as *f* (*N* (*τ, x*)), is given by:

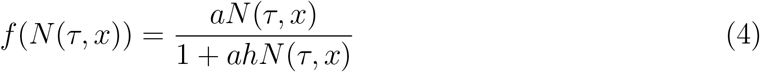

where parameters *a* and *h* represent the attack rate and handling time of the seed predators, respectively.

I assume that predators remove seeds by consumption but do not disperse them further. Therefore, the temporal dynamics of the seed density at a specific distance *x* and time *τ* after the dung beetle dispersal are described by:

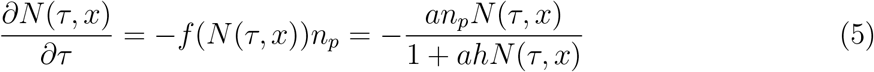

where *n_p_*is the density of seed predators. For simplicity, I assume that predators are uniformly distributed across space and time; thus, *n_p_*is a constant parameter independent of time *τ* and distance *x*.

I assume that the time elapsed from seed dispersal to germination, *T*, is identical across all seeds. The seed density at the time of germination, *N* (*T, x*), is obtained as follows:

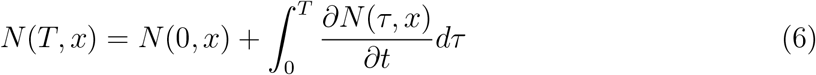

In Scenario A, predators are absent (*n_p_* = 0); therefore, the seed density at time *T* is equal to that at time 0 (*N* (*T, x*) = *N* (0, *x*)).

#### 2.1.4 Number of established seedlings

Seedlings compete for limited resources, including water, nutrients, space, and light (Coomes and Grubb 2000). I incorporate among-seedling competition through a negative density-dependent establishment probability, where the probability that a single seed at location *x* successfully establishes as a seedling decreases with the local density of surviving seeds, *N* (*T, x*). Thus, in a scenario governed solely by competition among seedlings, the establishment probability of a seed at location *x*, denoted by *S*(*N* (*T, x*)), is given by:

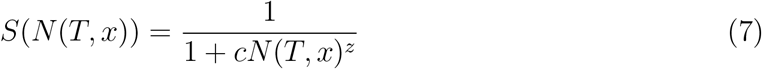

where *c* is the competition coefficient and *z* is a scaling parameter that determines how sharply the establishment probability declines with increasing seed density.

The total number of established seedlings in the entire focal habitat is calculated as the product of the total dung consumption by the dung beetle community and the number of seedlings established from a single dung pat. Consequently, the total number of established seedlings, *E*, is given by:

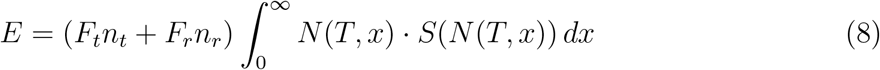

### 2.2 Simulation Methods

Based on the behavioral experiments of Manns et al. 2020, I parameterized the seed dispersal kernel for each dung beetle functional group. For the Gamma distribution of rollers, the parameters were set to yield a mean dispersal distance of 150 cm and a variance of 3600 cm², resulting in *k_r_* = 6.25 and *θ_r_* = 24.0. For tunnelers, the parameters of the Gamma distribution were set to *k_t_*= 1 and *θ_t_*= 10. All parameters used in this study are summarized in Table 1.

**Table 1:** Notation and parameters used.

| Notation. | Description. | Value. |
| --- | --- | --- |
| $n_t, n_r$ | Population sizes of tunnelers and rollers. | [0 - 100] |
| $s, 1 - s$ | Proportions of tunnelers and rollers | [0 - 1] |
| $K$ | Carrying capacity of dung beetles | [100] |
| $D_i(x)$ | Probability density function of dispersal distance [$i \in \{t, r\}$] | |
| $x$ | Distance from the dung pat. | [0 - 500] |
| $k_i$ | Shape parameter of the Gamma distribution [$i \in \{t, r\}$] | $[1, \frac{100}{36}]$ |
| $\theta_i$ | Scale parameter of the Gamma distribution [$i \in t, r$] | $[10, \frac{1}{36}]$ |
| $N(x)$ | Local density of dispersed seeds at distance $x$. | |
| $I$ | Total seed number in a unit of feces dispersed in one direction | [100] |
| $F_i$ | Amount of dung consumed by a single dung beetle [$i \in t, r$] | [1] |
| $E$ | Total number of established seedlings | |
| $S(N(x))$ | Seedling establishment probability | |
| $c$ | Competition coefficient among seeds | [0 - 4 ] |
| $z$ | Scaling parameter of density-dependent competition | [1] |
| $\tau$ | Time from dispersal by dung beetles | [1] |
| $T$ | Time from dispersal to germination | [10] |
| $f(N(x))$ | Predation amount | [1] |
| $h$ | Handling time of predators | [2, 4, 6 ] |
| $a$ | Attack rate of predators | [1] |
| $P(N(T, x))$ | Probability of seed survival after predation. | |

Although the model theoretically considers dispersal distances from 0 to *∞*, I truncated the spatial domain (*x*) to a finite range of [0, 500] cm in the numerical simulations for computational tractability. Under this parameter set, the two functional groups of dung beetles scarcely dispersed seeds beyond 500 cm; therefore, this truncation did not qualitatively affect the results (see Fig. S1). The total number of established seedlings (*E*) was calculated by numerically integrating Eq. (8) over *x*.

To quantify how beetle community composition affects plant recruitment, I varied the proportion of tunnelers (*s*) from 0 (rollers-only) to 1 (tunnelers-only) in increments of 0.025. I calculated the relative change in the number of established seedlings in the presence of tunnelers compared to a rollers-only community (*s* = 0), denoted as Δ*E*:

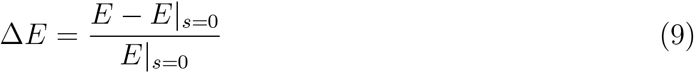

Positive values of Δ*E* indicate that tunneler presence enhances recruitment; negative values indicate reduced recruitment.

In Scenario B, I derived an implicit analytical solution for the temporal dynamics of seed density via separation of variables (Eqs. (3), (5), (6)). I then solved this implicit equation numerically using Brent’s method (Virtanen et al. 2020; for the derivation, see Supplement S1). Let *P* (*N* (*T, x*)) denote the survival probability from predation until germination, given by *P* (*N* (*T, x*)) = *N* (*T, x*)*/N* (0, *x*).

To investigate how post-dispersal processes interfere with seedling establishment, I systematically varied the intensity of competition and predation. For the former, I varied the competition coefficient (*c*) from 0 (no competition effect) to 4 (in increments of 0.04). For the latter, which is incorporated only in Scenario B, I simulated three distinct levels of predation pressure by varying the handling time (*h*): a short handling time indicating high predation pressure (*h* = 2), an intermediate handling time and medium predation pressure (*h* = 4), and a long handling time and weak predation pressure (*h* = 6). All numerical analyses were performed in Python (version 3.13.9) using the NumPy, SciPy and matplotlib packages. The Python code used for all analyses and to generate the results is publicly available on GitHub (https://anonymous.4open.science/r/Dungbeetle Seed dispersal Model-B07C/README.md).

## 3 Result

### Scenario A: Competition-driven establishment

In Scenario A, which considered only intraspecific competition among seedlings (*n_p_* = 0) as a post-dispersal process, Δ*E* showed a non-monotonic change with increasing proportion of tunneler species (Fig. 2a). Specifically, at low tunneler proportions (*s ≈* 0 *−* 0.25), total seedling establishment increased compared to a rollers-only community (Δ*E >* 0).

**Figure 2:**
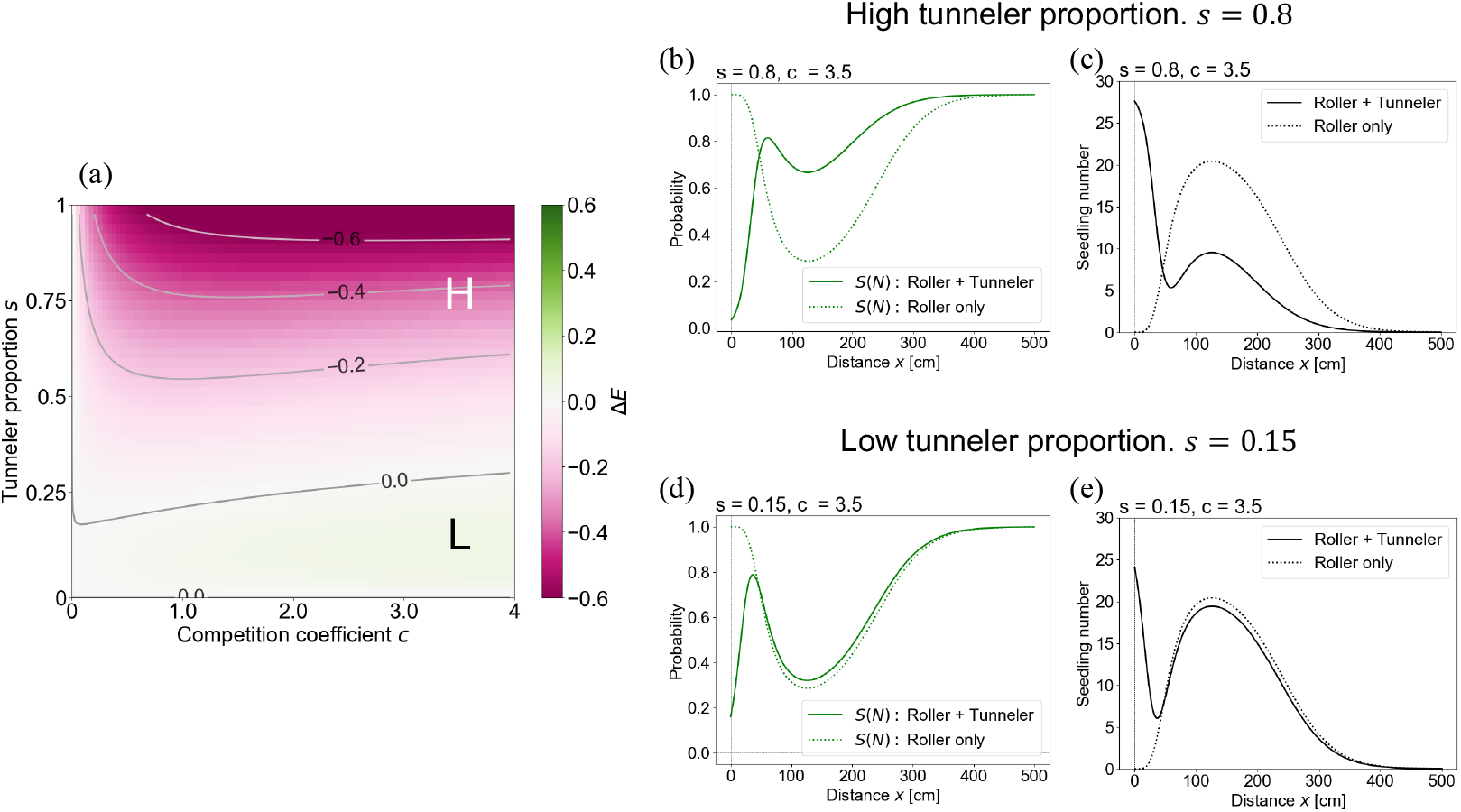
Effect of community composition on seedling establishment in Scenario A. (a) The relative difference in the total number of established seedlings (Δ*E*) under the competition-only scenario (Scenario A). Green and magenta shading represent an increase and decrease, respectively, relative to a rollers-only community (*s* = 0). Labels H and L indicate the parameter sets used in panels (b,c) and (d,e), corresponding to high and low tunneler proportions, respectively. (b), (d) Spatial distribution of the established local seedling number. (c), (e) The seedling establishment probability. The solid lines represent a mixed community of rollers and tunnelers, and the dotted lines represent a rollers-only community.

To examine the spatial distribution of established seedlings and their establishment probabilities (*S*(*N* (*T, x*))), we compared two parameter sets: *s* = 0.8, *c* = 3.5 (negative Δ*E*, denoted as H in Fig. 2a) and *s* = 0.15, *c* = 3.5 (positive Δ*E*, denoted as L in Fig. 2a). When Δ*E* was negative, although competition among roller-dispersed seeds was mitigated (Fig. 2b), the number of established seedlings was significantly lower compared to the rollers-only community (Fig. 2c). In contrast, when Δ*E* was positive, the seedling establishment probability of roller-dispersed seeds was nearly identical to that of the rollers-only community (Fig. 2d), and the number of established tunneler-dispersed seedlings more than compensated for the lower number of roller-dispersed seedlings (Fig. 2e). The competition coefficient *c* had minimal impact on Δ*E* except at very low values combined with high tunneler proportions.

A comparing of three values of the scaling parameter for competition, *z*, revealed that a higher *z* resulted in a more negative value of Δ*E*; namely, the number of established seedlings was lower than that of the rollers-only community (Fig. S2). Regarding the initial number of seeds per dung pat, *I*, the parameter region corresponding to the sign of Δ*E* shifted depending on *I* (Fig. S3). A smaller initial number of seeds caused an expansion the positive region of Δ*E*, while a larger number caused an expansion of the negative region. In other words, a higher initial number of seeds led to a greater decline in the number of established seedlings in the mixed community compared to the rollers-only community.

### Scenario B: Combined effects of competition and predation

In Scenario B, which considered seed predation, the difference in the number of established seedlings (Δ*E*) exhibited non-monotonic change with respect to the tunneler proportion, *s* (Fig. 3). Specifically, when the handling time of seed predators was short (*h* = 2), the total seedling establishment increased (Δ*E >* 0) with an increasing tunneler proportion *s* (Fig. 3a), in contrast to Scenario A. Notably, the total number of established seedlings was lowest at a high roller proportion of 0.9 (i.e., *s ≈* 0.1), which was also opposite to the pattern observed in Scenario A (Fig. 2a).

**Figure 3:**
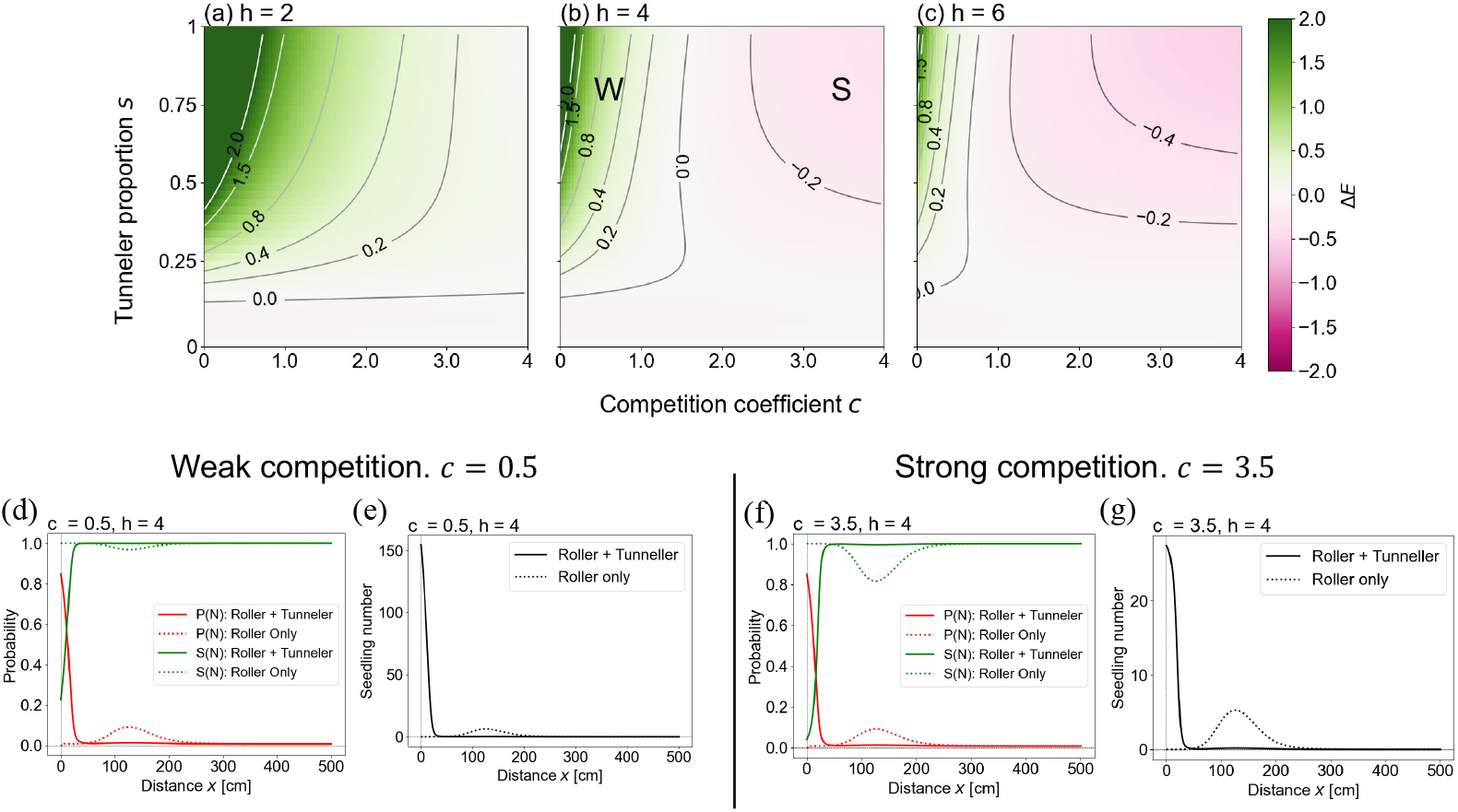
Effect of community composition on seedling establishment under both competition and predation (Scenario B). (a)–(c) The relative difference in the total number of established seedlings (Δ*E*) under varying handling times: (a) short (*h* = 2), (b) intermediate (*h* = 4), and (c) long (*h* = 6). Green and magenta shading represent an increase and decrease, respectively, relative to a rollers-only community (*s* = 0). Labels W and S in panel (b) indicate the parameter sets used for the spatial profiles in panels (d, e) and (f, g), corresponding to weak (*c* = 0.5) and strong (*c* = 3.5) competition under medium handling time, respectively. (d), (f) Spatial distribution of the established local seedling number. (e), (g) The underlying ecological probabilities. The green lines represent the seedling establishment probability under competition, *S*(*N*), and the red lines represent the survival probability from predation, *P* (*N*). The solid lines represent a mixed community of rollers and tunnelers, and the dotted lines represent a rollers-only community.

At an intermediate handling time (*h* = 4; Fig. 3b), the effect of tunnelers depended critically on the competition coefficient *c*. A comparison of two representative parameter sets sharing the same tunneler proportion (*s* = 0.8) but yielding opposite signs for Δ*E* was conducted to examine the spatial distribution of the number of established seedlings, their establishment probabilities (*S*(*N* (*T, x*))), and their survival probabilities (*P* (*N* (*T, x*))). Under weak competition (*c* = 0.5), the establishment probabilities of tunneler-dispersed seeds were high and those in the rollers-only community were also high (Fig. 3d). Consequently, the number of established tunneler-dispersed seedlings more than compensated for the reduction in roller-dispersed seedlings (Fig. 3e); thus Δ*E* was positive. In contrast, under strong competition (*c* = 3.5), the establishment probabilities of tunneler-dispersed seeds were low (Fig. 3f); therefore, the number of established tunneler-dispersed seedlings was lower than that in the rollers-only community (Fig. 3g).

Under a long handling time for predators (*h* = 6; Fig. 3c), total seedling establishment became increasingly lower than that in the rollers-only community (Δ*E ≤* 0) as the competition coefficient (*c*) increased, except at very low values ( *c ≤* 0.8). This decline was more pronounced than that observed under an intermediate handling time.

The scaling parameter for competition, *z*, and the initial number of seeds per dung pat, *I*, similarly influenced the response of total seedling establishment to dung beetle community composition (Figs. S4, S5). Specifically, at low values of *z* or *I*, the parameter region yielding a positive Δ*E* was dominant, whereas higher values of these parameters expanded the region where Δ*E <* 0.

## 4 Discussion

### 4.1 Context-dependent effects of dung beetle community composition

This model reveals that the optimal dung beetle community for maximizing seedling establishment is fundamentally context-dependent, strictly governed by the balance between density-dependent competition and seed predation. Under a purely competition-driven scenario (Scenario A), niche complementarity maximizes total establishment when a small proportion of tunnelers coexists with a majority of rollers. However, the presence of seed predators (Scenario B) completely reverses this pattern. Specifically, high predation pressure shifts the advantage toward tunneler-dominated communities, as local seed concentration induces predator satiation. At intermediate predation pressures, the optimal community depends on competition strength, whereas weak predation restores the superiority of rollers because density-dependent mortality from local crowding exceeds predation-induced mortality. Ultimately, these findings demonstrate that the ecological value of spatial dispersion by rollers versus physical refuge provided by tunnelers shifts dynamically in response to prevailing post-dispersal mortality rates.

In the absence of seed predation, niche complementarity maximizes seedling establishment (Fig. 2a). Under a purely competition-driven scenario, total establishment peaks when a small proportion of tunnelers coexists with a majority of rollers. Tunnelers bury a small fraction of seeds in a distinct spatial niche―the area immediately beneath or near the dung pat―which is largely unutilized by rollers. This spatial separation results in a high establishment probability for tunneler-dispersed seeds (Fig. 2e), more than compensating for the reduction in roller-dispersed seedlings (Fig. 2d). However, as the tunneler proportion increases beyond this optimum, intensified local competition among tunneler-dispersed seeds decreases overall establishment. Consequently, in ecosystems where rollers naturally constitute a large proportion of the community, such as tropical savannas (Kunz and KRELL 2011), an increase in tunnelers would likely decrease total seedling establishment, assuming negligible predation pressure.

Alongside community composition, predation pressure critically dictates seedling establishment. Under high predation pressure, a higher tunneler proportion maximizes total establishment, as local seed concentration by tunnelers induces predator satiation and increases individual seed survival (Fig. 3a). Conversely, with long handling times and weak predation (Fig. 3c), roller-dispersed seeds easily survive, enabling the rollers-only community to achieve higher establishment. By extension, Scenario A, which considered only seedling competition, can be interpreted as the predator handling time *h* being exceptionally long. Furthermore, under intermediate predation (Fig. 3b), outcomes depend on competition strength. Weak competition results in higher total establishment in tunneler-dominated communities (Fig. 3e), because established seedlings from tunnelers outnumbere roller-dispersed ones, which have high predation rates (Fig. 3d). Conversely, strong competition keeps the establishment probability of tunneler-dispersed seeds low (Fig. 3f), leading to higher total establishment in the rollers-only community (Fig. 3g). Notably, across all cases with predation, communities consisting of a small proportion of tunnelers and a large proportion of rollers exhibit lower total seedling establishment than the rollers-only community. This reduction occurs because seeds are distributed most sparsely in this condition, resulting in high mortality of the dispersed seeds due to predation.

### 4.2 Ecological implications of anthropogenic shifts in community structure

Shifts in dung beetle community structure driven by anthropogenic disturbance significantly alter total seedling establishment. In Madagascar, numerous dung beetle species went extinct due to deforestation and the loss of large native mammals (Hanski et al. 2007). Furthermore, veterinary antiparasitic drugs such as ivermectin cause sensory and locomotor disorders (Verdú et al. 2015), increase mortality (Martínez et al. 2017), reduce larval emergence rates (González-Tokman et al. 2017), and consequently reshape community structure (Sands and Wall 2018) and ecosystem function (Verdú et al. 2018). In particular, roller dung beetles, often having narrower dietary breadths, are highly vulnerable to population declines (Akashi et al. 2024; Carpaneto et al. 2007; Salomao et al. 2019). For example, while tunneler dung beetles can exploit non-native cattle dung, certain roller species in Mexico fail to utilize it (Amézquita and Favila 2010). When these disturbances decrease the proportion of rollers and increase that of tunnelers, this model predicts context-dependent outcomes: under competition-dominant conditions, total seedling establishment decreases, whereas under strong predation pressure, the opposite outcome occurs. Therefore, to accurately evaluate the impact of anthropogenic disturbance on secondary seed dispersal, future studies must account not only for which dung beetle guilds increase or decrease, but also for environmental conditions, including plant establishment traits and local predation pressure.

The intentional introduction of dung beetles is a widespread practice globally to secure substantial economic benefits from ecological services like dung decomposition and plant growth enhancement (Losey and Vaughan 2006). Consequently, highly efficient dwellers and tunnelers are frequently introduced, along with occasional introductions of African rollers intended to promote long-distance secondary seed dispersal (Pokhrel et al. 2021). However, results indicate that introducing either rollers or tunnelers can unexpectedly decrease total seedling establishment, driven by the interactions between seed predators and the native community composition. Furthermore, because native plants are evolutionarily adapted to the specific burial depths provided by native dung beetles (Koike et al. 2012), non-native dung beetles may bury seeds at suboptimal depths. In such cases, the introduction of non-native dung beetles could further decrease total seedling establishment. Ultimately, the artificial introduction of dung beetles and resulting shifts in community structure may inhibit pasture grass renewal, thereby incurring unexpected economic costs.

Seed burial by dung beetles is known to reduce predation mortality (Andresen and Levey 2004), offering an alternative mechanism for the high survival of tunneler-dispersed seeds beyond the assumed Type II functional response. While this model assumes a spatially uniform predator distribution, natural predators utilize olfactory cues to forage (Parker et al. 2023). If predator density *n_p_* acts as a function of local seed density *N* (*x*), density-dependent foraging would increase mortality in highly concentrated patches. Under such conditions, roller-dominated communities could maximize total establishment even under high predation pressure (Fig. 3a). Conversely, high local seed density can accelerate germination in certain plants (Orrock and Christopher 2010). In this scenario, the time to germination *T* becomes a function of initial local seed density *N* (0, *x*), meaning dense aggregations could shorten exposure time and reduce mortality. Therefore, accurately evaluating post-dispersal mortality in empirical systems strictly requires accounting for both plant germination traits and predator foraging behaviors.

Beyond predation, other ecological mechanisms can explain the increased seedling establishment in highly aggregated seed patches. While this model explicitly formulates the dilution effect against predation pressure, positive density dependence can also emerge from facilitative interactions, such as nurse plant effects, local soil modification, chemical resource sharing, and mycorrhizal networks. Specifically, high seed density enables the efficient sharing of mycorrhizal fungi. Ectomycorrhizal plants are known to promote conspecific seedling establishment, whereas arbuscular mycorrhizal (AM) plants inhibit it (Kadowaki et al. 2018). However, AM plants have adaptively evolved for endozoochory (Yamawo and Ohno 2024). Therefore, for AM plants, local seed aggregation and fungal sharing act to intensify intraspecific competition. Consequently, AM plants would maximize seedling establishment when dispersed by rollers, whereas ectomycorrhizal plants would achieve higher establishment when dispersed by tunnelers.

### 4.3 Limitations and Future research questions

In natural ecosystems, secondary seed dispersal by dung beetles inherently operates within two- and three-dimensional spatial structures. This model assumes that dung pats are sufficiently isolated and therefore does not explicitly incorporate the spatial distribution of dung or the overlap of relocated seeds. The spatial distribution of seeds (seed shadow) following endozoochory is determined by the interaction between mammalian defecation patterns and secondary seed dispersal by dung beetles. Therefore, if mammals prefer certain defecation spots (Buesching et al. 2016; Miguel et al. 2018; Quirós-Navarro et al. 2026) or synchronize their defecation timing (P. Rees 1983), the resulting dung aggregation can cause even roller-dispersed seeds to become spatially clustered. While this study formulated seed dispersal as a one-dimensional process, rollers are known to transport brood balls in linear trajectories using celestial cues to minimize intense competition at the dung patch (Dacke et al. 2013a,b). Considering a two-dimensional space, local seed density would be described more complexly. Moreover, if roller dung beetles exhibit a preference for burial microhabitats, this spatial overlap of seeds will become even more pronounced (see Baird et al. 2010). Furthermore, this model does not incorporate the vertical dimension of dung burial depth. Generally, tunnelers bury brood balls deep in the soil, whereas rollers bury them shallowly near the surface (Halffter and Edmonds 1982). When seeds are buried beyond a certain depth, the establishment probability decreases (Andresen and Levey 2004). Notably, some tunneler dung beetles bury seeds deeper than the critical depth for germination (Koike et al. 2012). Seeds buried at such depths do not germinate immediately, but instead enter the soil seed bank. Therefore, future theoretical models must evolve toward spatially explicit approaches that integrate both mammalian dung distribution and the two-dimensional dispersal trajectories of dung beetles (see Yin and Zinn-Björkman 2020) while also formulating guild- and species-specific burial depths.

In this model, I did not explicitly distinguish between incorporation into the seed bank (dormancy) and mortality due to germination failure. Although high seed density can some-times promote seed dormancy (Murray 1998), seeds ingested by animals typically exhibit enhanced germination due to a combination of mechanisms (Razanamandranto et al. 2004). These mechanisms include the removal of pulp and associated germination inhibitors (Yagihashi et al. 2000), scarification of the seed coat (H. Ishikawa 2010; Janzen 1981), and exposure to gastric acids in the digestive tract (Gardener et al. 1993). Therefore, ingested seeds dispersed by dung beetles are likely to have a lower probability of entering dormancy compared to uningested seeds. Furthermore, short-lived plants tend to rely more heavily on seed dormancy to avoid extinction risk, acting as a bet-hedging strategy (M. Rees 1996). Consequently, this modeling framework may be more applicable to long-lived plants (e.g., perennial herbs or trees). A key limitation of this study, therefore, is its inability to account for the dynamics of annuals. In addition to the density-dependent mortality addressed in this study, explicitly incorporating processes such as seed dormancy would enable future models to evaluate seed dispersal and seedling establishment specifically for these short-lived taxa. This model does not explicitly account for the specific species of mammalian defecators.

The abundance and species richness of seeds within a dung pat vary significantly depending on the defecator’s functional guild, such as frugivores versus herbivores. Specifically, a single dung pat from a frugivore generally exhibits a higher species richness of fruit-bearing plant seeds, whereas herbivore dung contains a significantly greater absolute abundance of herbaceous plant seeds (Karimi et al. 2020; Perea et al. 2013). Even among herbivores, contributions to endozoochory vary: larger mammals consume greater quantities of seeds (Shiferaw et al. 2004) and are less likely to destroy them during ingestion and digestion (Miller 1995; Neto et al. 1987). The number of seeds contained within a dung pat leads to divergent outcomes for seedling establishment (Figs. S3, S5). Furthermore, in environments where multiple dung types coexist, the dung preference of beetles becomes a critical factor. Although this study assumed uniform preference and consumption capacity among dung beetle guilds (*F_r_*= *F_t_*), dung beetles in natural ecosystems exhibit distinct preferences and utilization efficiencies for different mammalian dung types (Bogoni and Hernández 2014; Onodera et al. 2022; Whipple and Hoback 2012). Therefore, to accurately evaluate the secondary seed dispersal function of dung beetles, future studies must explicitly incorporate the identities of mammalian endozoochorous dispersers, alongside the corresponding dung preferences and utilization capacities of the local dung beetle community.

