## Supplement S1 for "How does the community structure of dung beetles affect ecosystem functions? Modeling of secondary seed dispersal and seedling establishment"

### 1 Derivation.

In this derivation, I analytically calculate the seed density surviving predation, $N(T, x)$. From Eq.7, the temporal dynamics of the seed density at a specific distance $x$ with time $\tau$ after dispersal are described by:

$$\frac{\partial N(\tau, x)}{\partial \tau} = -\frac{an_p N(\tau, x)}{1 + ahN(\tau, x)} \quad (1)$$

where $n_p$ is the density of seed predators, $a$ is the attack rate, and $h$ is the handling time. To find the remaining seed density at the time of germination ($\tau = T$), I solve this equation analytically by separating the variables. Treating $x$ as a constant spatial parameter, I rearrange the equation as follows:

$$\frac{1 + ahN(\tau, x)}{N(\tau, x)} dN(\tau, x) = -an_p d\tau \quad (2)$$

$$\left( \frac{1}{N(\tau, x)} + ah \right) dN(\tau, x) = -an_p d\tau \quad (3)$$

Next, I integrate both sides over the time interval from dispersal ($\tau = 0$) to germination ($\tau = T$). The corresponding initial and final seed densities are $N(0, x)$ and $N(T, x)$, respectively:

$$\int_{N(0, x)}^{N(T, x)} \left( \frac{1}{N} + ah \right) dN = \int_0^T -an_p d\tau \quad (4)$$

Solving the integrals yields:

$$[\ln(N) + ahN]_{N(0,x)}^{N(T,x)} = [-an_p\tau]_0^T \quad (5)$$

$$(\ln(N(T, x)) + ahN(T, x)) - (\ln(N(0, x)) + ahN(0, x)) = -an_pT \quad (6)$$

By combining the logarithmic terms, I obtain the implicit equation for $N(T, x)$:

$$\ln\left(\frac{N(T, x)}{N(0, x)}\right) + ah(N(T, x) - N(0, x)) = -an_pT \quad (7)$$

Because this implicit equation cannot be solved algebraically for $N(T, x)$ in terms of elementary functions, I determined $N(T, x)$ numerically in the simulations. Specifically, I computed the roots of this non-linear equation using Brent’s method, implemented via the `brentq` function in the ‘`scipy.optimize`’ module in Python (Virtanen et al. 2020). Once $N(T, x)$ is numerically obtained, the survival probability from predation is calculated as $P(N(0, x)) = \frac{N(T, x)}{N(0, x)}$.
