## Supplementary material for "How does the community structure of dung beetles affect ecosystem functions? Modeling of secondary seed dispersal and seedling establishment": Figure S

### 1 Supplementary Figures

Figure S1

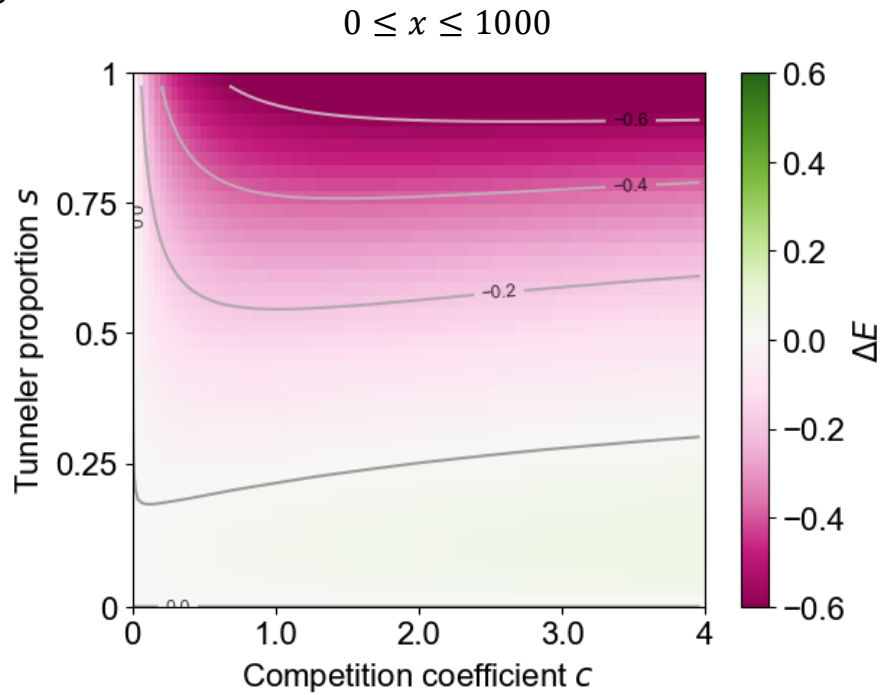

Fig. S1: Effect of community composition on seedling establishment in Scenario A when the upper limit of distance, $x$, is set to 1000 cm. The relative difference in the total number of established seedlings ($\Delta E$) under the competition-only scenario (Scenario A). Green and magenta shading represent an increase and decrease, respectively, relative to a rollers-only community ($s = 0$).

Figure S2

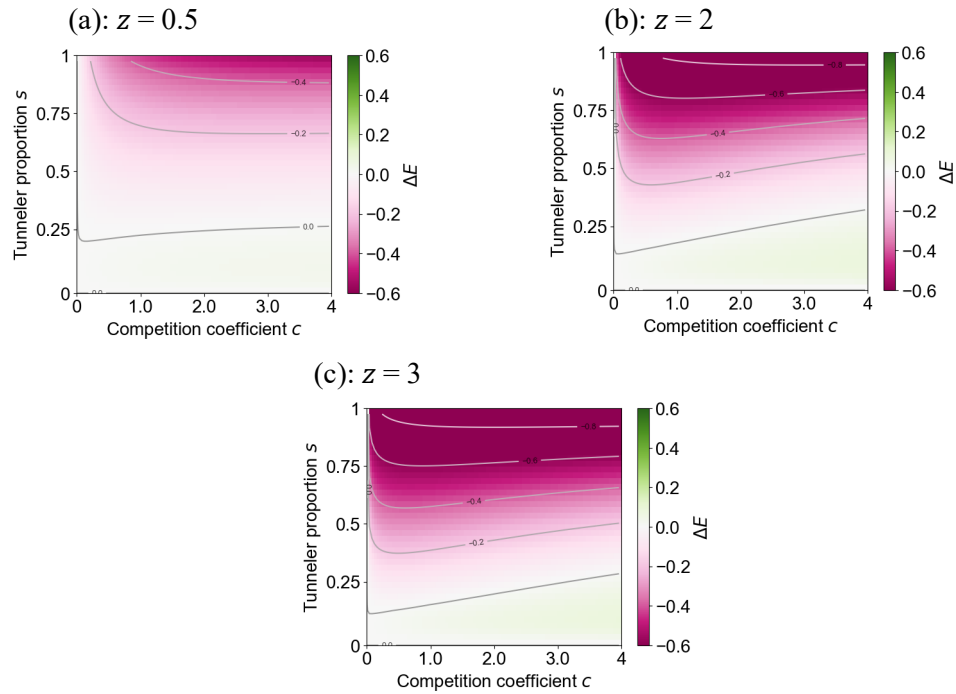

Fig. S2: Effect of the scaling parameter for competition, $z$, on seedling establishment in Scenario A. Panels show the relative difference in the total number of established seedlings ($\Delta E$) for (a) $z = 0.5$, (b) $z = 2$, and (c) $z = 3$. Green and magenta shading represent an increase and decrease, respectively, relative to a rollers-only community ($s = 0$).

Figure S3

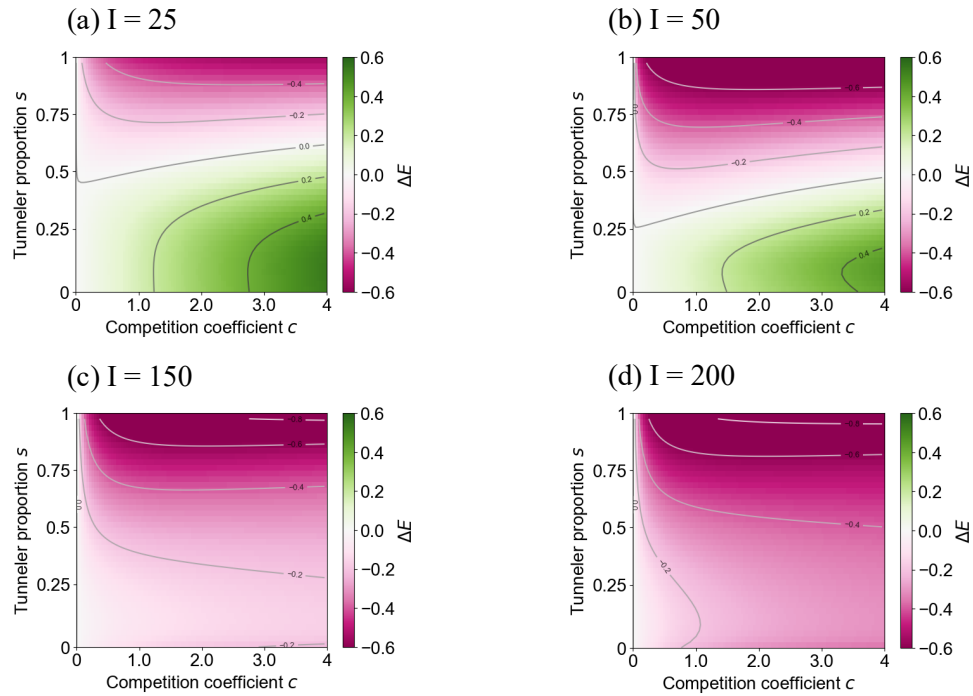

Fig. S3: Effect of initial number of seeds per dung pat, $I$, on seedling establishment in Scenario A. Panels show the relative difference of the total number of established seedlings ($\Delta E$) for (a) $I = 25$, (b) $I = 50$, (c) $I = 150$, and (d) $I = 200$. Green and magenta shading represent an increase and decrease, respectively, relative to a rollers-only community ($s = 0$).

Figure S4

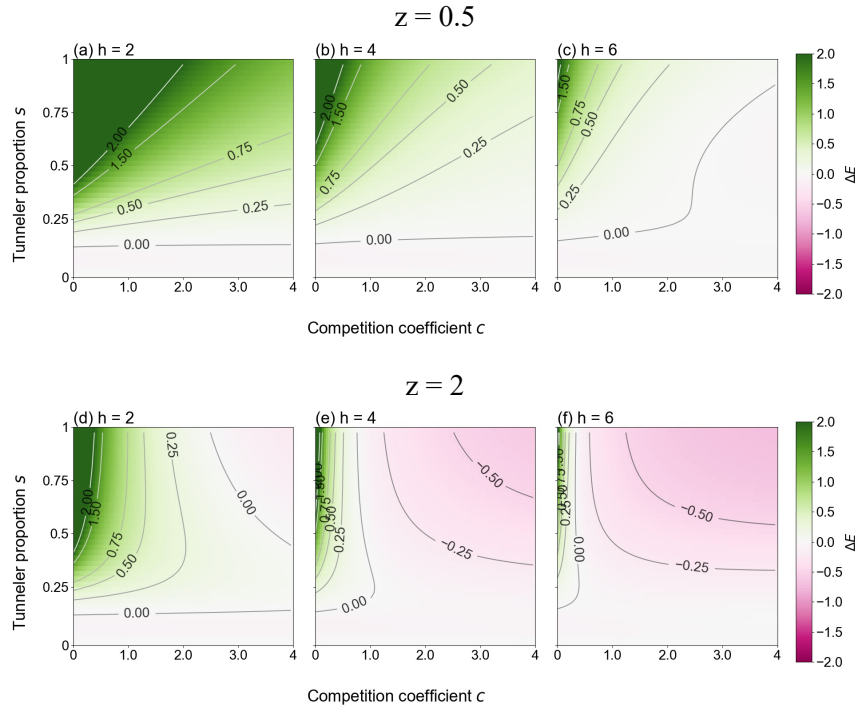

Fig. S4: Effect of the scaling parameter for competition, $z$, on seedling establishment under both competition and predation (Scenario B). Panels show the relative difference in the total number of established seedlings ($\Delta E$) under varying handling times: (a, d) short ($h = 2$), (b, e) intermediate ($h = 4$), and (c, f) long ($h = 6$). The scaling parameter is $z = 0.5$ for (a)–(c) and $z = 2$ for (d)–(f). Green and magenta shading represent an increase and decrease, respectively, relative to a rollers-only community ($s = 0$).

Figure S5

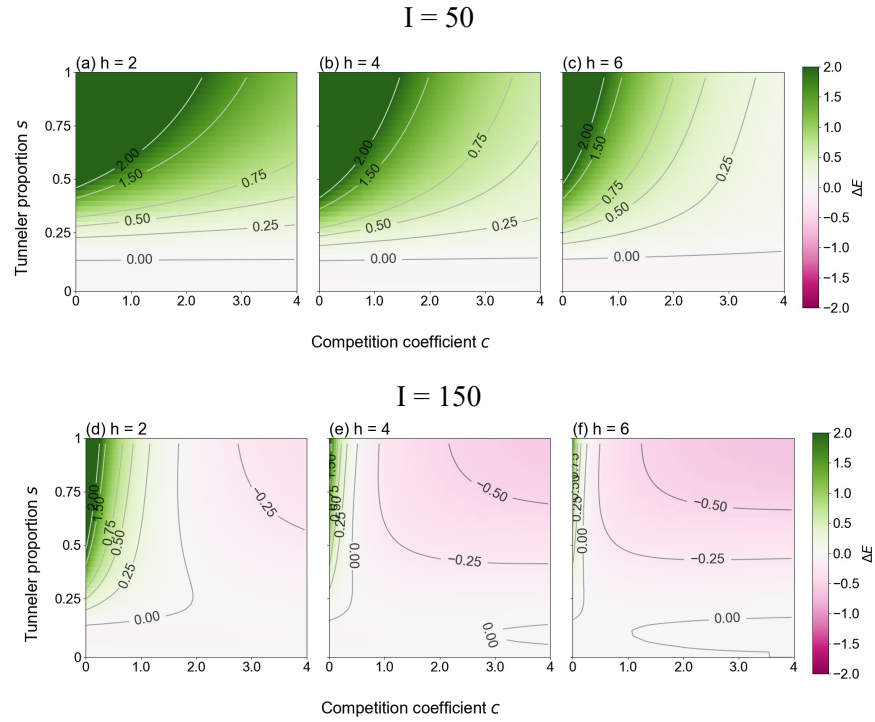

Fig. S5: Effect of the initial number of seeds per dung pat, $I$, on seedling establishment under both competition and predation (Scenario B). Panels show the relative difference in the total number of established seedlings ($\Delta E$) under varying handling times: (a, d) short ($h = 2$), (b, e) intermediate ($h = 4$), and (c, f) long ($h = 6$). The initial number of seeds is $I = 50$ for (a)–(c) and $I = 150$ for (d)–(f). Green and magenta shading represent an increase and decrease, respectively, relative to a rollers-only community ($s = 0$).
